# Optimization and validation of a quadruplex real-time PCR assay for the diagnosis of diphtheria

**DOI:** 10.1101/600270

**Authors:** Edgar Badell, Sophie Guillot, Marie Tulliez, Marine Pascal, Leonardo-Gabriel Panunzi, Samuel Rose, David Litt, Norman K. Fry, Sylvain Brisse

**Author notes:** **Corresponding author**: Sylvain Brisse. Biodiversity and Epidemiology of Bacterial Pathogens, Institut Pasteur, 25 rue du Docteur Roux, F-75724 Paris, France.

## Abstract

Diphtheria is caused by toxigenic strains of *Corynebacterium diphtheriae, Corynebacterium ulcerans* and *Corynebacterium pseudotuberculosis*. For diagnostic purposes, species identification and detection of toxigenic strains (diphtheria toxin (tox)-positive strains) is typically performed using end-point PCR. A faster quadruplex real-time PCR (qPCR) was recently developed (De Zoysa *et al*. J Med Microbiol. 2016 65(12):1521-1527). Here, we present an improvement of the quadruplex method, in which a 16S rRNA gene target was added as an internal processing control, providing confirmation of the presence of bacterial DNA in the assays. This improved qPCR method was validated using 36 bacterial isolates and 16 clinical samples. The method allows detection of the *tox* gene and distinguishing *C. diphtheriae* (including the newly described species *C. belfantii*) from *C. ulcerans* and *C. pseudotuberculosis*. Complete diagnostic specificity, sensitivity and experimental robustness of the method to temperature and reagent concentration variations were demonstrated. The lower limit of detection for *C. diphtheriae, C. ulcerans* and *tox* targets was 1.86 genome copies per 5 μL reaction volume. Finally, the method was successfully used on two distinct qPCR technologies (LightCycler 480, Roche Diagnostics and Rotor-Gene Q, Qiagen) and in two laboratories (Institut Pasteur, Paris, France and Public Health England – National Infection Service, London, UK). This work describes validation of the improved qPCR quadruplex method and supports its implementation for the biological diagnosis of diphtheria.

## Introduction

*Corynebacterium diphtheriae* is the main etiological agent of diphtheria, a once-common acute human infection classically affecting the upper respiratory tract and occasionally the skin. The severe manifestations of the disease are caused by the action of the diphtheria toxin, produced by some strains of *C. diphtheriae* which carry the *tox* gene. Strains of *Corynebacterium ulcerans* and more rarely *Corynebacterium pseudotuberculosis* can also be toxigenic *i.e*., be capable of secreting the toxin, and can cause infections in humans. The three species are phylogenetically related and we collectively define them as the *C. diphtheriae* species complex. Recently a subset of *C. diphtheriae* strains of one of the four biovars, Belfanti, were recognized as forming a novel species, *C. belfantii* (1). Although this novel species also belongs to the *C. diphtheriae* complex, *C. belfantii* strains generally do not carry the *tox* gene (1, 2).

Diphtheria is a well-controlled disease in countries with high vaccination coverage. However, the vaccine targets the toxin but does not prevent transmission of bacteria of the *C. diphtheriae* complex, and low coverage or discontinuation of vaccination can result in a rapid resurgence of diphtheria (3, 4). Further, *C. ulcerans* infections in humans have emerged recently and usually involve close contacts with animals, mainly domestic cats and dogs (5, 6). *C. pseudotuberculosis* is primarily a veterinary pathogen that infects ungulates such as sheep and goats (7), and the rare human infections with *C. pseudotuberculosis* are associated with occupational risk factors (8–10). Although rarely reported, *C. diphtheriae* can also infect animals such as cats, cows and horses (11–13). Identification of putative toxigenic corynebacteria at species level is classically performed by biochemical phenotypic methods (14–16) and more recently by matrix-assisted laser desorption/ionization time of flight (MALDI-TOF) mass spectrometry (17, 18). However, phenotypic methods require strain culture and isolation and are slow. In addition, the biochemical identification of *C. pseudotuberculosis* and its differentiation from other corynebacteria, especially *C. ulcerans*, can be difficult (10). Further, these methods cannot determine the toxigenic status of strains.

Determination of the potential toxigenic status of clinical isolates is the most critical aspect of diphtheria diagnosis, as it informs public health action and patient care, including possible treatment by administration of antitoxin. End-point PCR assays targeting the *tox* gene were developed in the 1990s (19–21) and are widely used to screen for the presence of potentially toxigenic strains directly from clinical samples or from bacterial cultures. Detection of the *tox* gene can also be combined with species identification PCR targets in multiplex assays (9). Because non-toxinogenic toxin-bearing (NTTB) isolates were described, the detection of the *tox* gene only provides presumption of toxigenicity, which can be confirmed using the Elek test (15).

Real-time PCR (qPCR, for quantitative PCR) presents the advantages of faster data collection than classical PCR, low contamination risks and high sensitivity. Several qPCR assays that target the *tox* gene have been described (22–25). Recently, a quadruplex qPCR assay for detection of the *tox* gene and identification of *C. diphtheriae, C. ulcerans* and *C. pseudotuberculosis* by targeting their RNA polymerase β-subunit (*rpoB*) gene sequences, was developed by De Zoysa *et al*. (26). For PCR diagnostic purposes, it is considered best practice to include process control(s) capable of detection both extraction failure and inhibition of PCR amplification (27). Whilst the De Zoysa *et al*. (21) method uses amplification of the green fluorescent protein (*gfp*) gene on control DNA to test for PCR inhibition, it does not include a control for extraction failure (*i.e*., one capable of detecting the presence/absence of bacterial DNA in the PCR assay).

Here, we aimed to address this limitation by replacing the *gfp* target gene by a universal fragment of the 16S rRNA (u-16S) gene sequence to serve as internal processing control. Further, we aimed to validate the improved qPCR assay directly on clinical specimens such as throat swabs and pseudomembrane biopsies. Additionally, we tested the characteristics of the modified quadruplex qPCR assay including specificity, sensitivity, reproducibility, experimental robustness and its implementation on distinct qPCR apparatuses and in separate laboratories.

## Materials and Methods

### Reference strains of the *Corynebacterium diphtheriae* complex

In experiments performed at the French National Reference Center, *C. diphtheriae* strain NCTC10648 (National Collection of Type Cultures, Public Health England, UK), which bears the *tox* gene (*tox+*), and *C. diphtheriae* strain NCTC10356, which is tox-negative (tox−), were used as positive and negative *tox* PCR controls, respectively, and as positive controls for *C. diphtheriae* identification. *Corynebacterium pseudotuberculosis* strain CIP102968^T^ and *Corynebacterium ulcerans* strain NCTC12077, which are both *tox*-, were used as controls for *C. pseudotuberculosis* and *C. ulcerans* identification, respectively. In the validation experiments at Public Health England, strains NCTC10648 and NCTC12077 were used as controls, as previously described (26).

### Clinical isolates, strains and specimens

Clinical isolates (n = 36), laboratory strains (n = 7) and specimens (n = 16) that had been previously characterized at the French National Reference Center for the Corynebacteria of the *diphtheriae* complex were included (**Table 1**).

**Table 1.** Strains, isolates and clinical samples analyzed.

| Reference strains | Species and tox status* | Universal 16S rRNA | *rpoB* C. diphtheriae | *rpoB* C. ulcerans/ C. pseudotuberculosis | tox gene | Toxin production (Elek test) | Conclusion |
| --- | --- | --- | --- | --- | --- | --- | --- |
| NCTC10356 | *C. diphtheriae* tox- | + | + | - | - | ND | *C. diphtheriae* tox- |
| NCTC10648 | *C. diphtheriae* tox+ | + | + | - | + | + | *C. diphtheriae* tox+ |
| FRC0043T | *C. belfantii* tox- | + | + | - | - | ND | *C. diphtheriae* tox- |
| NCTC12077 | *C. ulcerans* tox- | + | - | + | - | ND | *C. ulcerans* tox- |
| CIP102968 | *C. pseudotuberculosis* tox- | + | - | + | - | ND | *C. pseudotuberculosis* tox- |
| CIP A95 | *C. pseudotuberculosis* tox- | + | - | + | - | ND | *C. pseudotuberculosis* tox- |
| CIP 52.103 | *C. pseudotuberculosis* tox- | + | - | + | - | ND | *C. pseudotuberculosis* tox- |
| CIP 52.104 | *C. pseudotuberculosis* tox- | + | - | + | - | ND | *C. pseudotuberculosis* tox- |
| CIP 52.97 | *C. pseudotuberculosis* tox- | + | - | + | - | ND | *C. pseudotuberculosis* tox- |
| CIP 59.46 | *C. pseudotuberculosis* tox- | + | - | + | - | ND | *C. pseudotuberculosis* tox- |
| NCTC764 | *Corynebacterium striatum* | + | - | - | - | ND | Non *diphtheriae* complex |
| **Isolates** |  |  |  |  |  |  |  |
| 00-0744 | *C. belfantii* tox- | + | + | - | - | ND | *C. diphtheriae* tox- |
| 05-3187 | *C. belfantii* tox- | + | + | - | - | ND | *C. diphtheriae* tox- |
| 06-4305 | *C. belfantii* tox- | + | + | - | - | ND | *C. diphtheriae* tox- |
| FRC0074 | *C. belfantii* tox- | + | + | - | - | ND | *C. diphtheriae* tox- |
| FRC0223 | *C. belfantii* tox- | + | + | - | - | ND | *C. diphtheriae* tox- |
| FRC0250 | *C. belfantii* tox- | + | + | - | - | ND | *C. diphtheriae* tox- |
| FRC0301 | *C. belfantii* tox- | + | + | - | - | ND | *C. diphtheriae* tox- |
| FRC0566 | *C. diphtheriae* tox+ | + | + | - | + | + | *C. diphtheriae* tox+ |
| FRC0568 | *C. diphtheriae* tox- | + | + | - | - | ND | *C. diphtheriae* tox- |
| FRC0570 | *C. diphtheriae* tox- | + | + | - | - | ND | *C. diphtheriae* tox- |
| FRC0018 | *C. diphtheriae* tox+ | + | + | - | + | + | *C. diphtheriae* tox+ |
| FRC0076 | *C. diphtheriae* tox+ | + | + | - | + | - | *C. diphtheriae* tox+ |
| FRC0101 | *C. diphtheriae tox+* | + | + | - | + | - | *C. diphtheriae tox+* |
| FRC0114 | *C. diphtheriae tox+* | + | + | - | + | - | *C. diphtheriae tox+* |
| FRC0365 | *C. diphtheriae tox+* | + | + | - | + | - | *C. diphtheriae tox+* |
| FRC0011 | *C. ulcerans tox-* | + | - | + | - | ND | *C. ulcerans tox-* |
| FRC0012 | *C. ulcerans tox-* | + | - | + | - | ND | *C. ulcerans tox-* |
| FRC0042a | *C. ulcerans tox+* | + | - | + | + | + | *C. ulcerans tox+* |
| FRC0058 | *C. ulcerans tox+* | + | - | + | + | - | *C. ulcerans tox+* |
| FRC0187 | *C. ulcerans tox+* | + | + | - | + | - | *C. ulcerans tox+* |
| FRC0567 | *C. ulcerans tox+* | + | - | + | + | + | *C. ulcerans tox+* |
| FRC0569 | *C. ulcerans tox-* | + | - | + | - | ND | *C. ulcerans tox-* |
| 05-770 | *C. ulcerans tox-* | + | - | + | - | ND | *C. ulcerans tox-* |
| 05-146 | *C. ulcerans tox-* | + | - | + | - | ND | *C. ulcerans tox-* |
| UFBA C231 | *C. pseudotuberculosis tox-* | + | - | + | - | ND | *C. pseudotuberculosis tox-* |
| UFBA C232 pld- | *C. pseudotuberculosis tox-* | + | - | + | - | ND | *C. pseudotuberculosis tox-* |
| FRC0041 | *C. pseudotuberculosis tox-* | + | - | + | - | ND | *C. pseudotuberculosis tox-* |
| FRC0186 | *C. pseudotuberculosis tox-* | + | - | + | - | ND | *C. pseudotuberculosis tox-* |
| FRC0386 | *Corynebacterium amycolatum* | + | - | - | - | ND | Non *diphtheriae* complex |
| FRC0539 | *Corynebacterium aurimucosum* | + | - | - | - | ND | Non *diphtheriae* complex |
| FRC0388 | *Enterobacter aerogenes* | + | - | - | - | ND | Non *diphtheriae* complex |
| FRC0392 | *Enterococcus faecalis* | + | - | - | - | ND | Non *diphtheriae* complex |
| FRC0413 | *Propionibacterium avidum* | + | - | - | - | ND | Non *diphtheriae* complex |
| FRC0427 | *Bacillus clausii* | + | - | - | - | ND | Non *diphtheriae* complex |
| FRC0428 | *Streptococcus pyogenes* | + | - | - | - | ND | Non *diphtheriae* complex |
| FRC0572 | *Neisseria subflava* | + | - | - | - | ND | Non *diphtheriae* complex |
| Samples |  |  |  |  |  |  |  |
| FRC0540 | Nasopharyngeal aspiration/ kit DNeasy | + | - | - | - | ND | Non *diphtheriae* complex |
| FRC0058-12N | Nose swab/ boiling | + | - | - | - | ND | Non *diphtheriae* complex |
| FRC0058-15N | Nose swab/ boiling | + | - | - | - | ND | Non *diphtheriae* complex |
| FRC0058-12T | Throat swab/ boiling | + | - | - | - | ND | Non *diphtheriae* complex |
| FRC0058-15T | Throat swab/ boiling | + | - | - | - | ND | Non *diphtheriae* complex |
| FRC0058-06 | Throat swab/ boiling | + | - | - | - | ND | Non *diphtheriae* complex |
| FRC0058-07 | Throat swab/ boiling | + | - | - | - | ND | Non *diphtheriae* complex |
| FRC0058-08 | Throat swab/ boiling | + | - | - | - | ND | Non *diphtheriae* complex |
| FRC0058-09 | Throat swab/ boiling | + | - | - | - | ND | Non *diphtheriae* complex |
| FRC0541 | Throat swab/ kit Dneasy | + | - | - | - | ND | Non *diphtheriae* complex |
| FRC0060 | Pseudomembrane/ kit Dneasy | + | - | - | - | ND | Non *diphtheriae* complex |
| FRC0064 | Pseudomembrane/ kit Dneasy | + | - | - | - | ND | Non *diphtheriae* complex |
| FRC0018 | Pseudomembrane/ kit Dneasy | + | + | - | + | ND | *C. diphtheriae* tox+ |
|  | Pseudomembrane/ boiling | + | + | - | + | ND | *C. diphtheriae* tox+ |
| FRC0042a | Pseudomembrane/ kit Dneasy | + | - | + | + | ND | *C. ulcerans* tox+ |
|  | Pseudomembrane/ boiling | + | - | + | + | ND | *C. ulcerans* tox+ |
| FRC0051 | Pseudomembrane/ kit Dneasy | + | - | - | - | ND | Non *diphtheriae* complex |
|  | Pseudomembrane/ boiling | + | - | - | - | ND | Non *diphtheriae* complex |
| FRC0058 | Pseudomembrane/ kit Dneasy | + | - | + | + | ND | *C. ulcerans* tox+ |
|  | Pseudomembrane/ boiling | + | - | + | + | ND | *C. ulcerans* tox+ |

\* Initial species identification was defined by end point PCR and/or MALDI-TOF and the *tox* status was defined by end point PCR.
FRC0058-12N; FRC0058-12T; FRC0058-15N; FRC0058-15T are swabs samples from nose (N) or throat (T) of
contacts of patient FRC0058. For contacts from FRC0058-06 to FRC0058-09 only throat samples were taken.
ND: Not done

### DNA extraction by the boiling method

DNA extraction was performed as follows. For bacterial strains, the method described by De Zoysa *et al*. (21) was used. For clinical swab material, swabs were introduced into a DNA/DNase/RNase free 1.5 ml Eppendorf Biopur tube (Cat. N° 0030 121.589, Eppendorf, Germany) containing 500 μl of nuclease free water (Cat. N°. P119C/Promega/U.S.A). The upper part of the swabs was cut using sterile scissors to allow closing of the tube. The tubes were vortexed thoroughly and placed in a preheated heating block at 100°C for 15 min. The swabs were then removed from tubes using sterile forceps, and the tubes centrifuged for 1 min at 13,000 g to pellet cell debris. For tissue samples, a piece of ca. 1 square cm of pseudomembrane was cut using sterile dissection forceps and scissors and introduced into a DNase/RNase free 1.5 ml Eppendorf tube as described above. The sample was ground using a sterile mini-grinder until a homogeneous suspension was obtained. The tubes were placed in a preheated heating block at 100°C for 15 min and then vortexed and centrifuged for 1 min at 13,000 g to pellet cell debris. The collected supernatant was used as template DNA for the PCR. A similar tube containing only 500 μl of nuclease-free water was included as no template control (NTC) for each extraction. Following the final centrifugation step for each sample type the supernatant was transferred to a new tube and used as template DNA for the PCR.

### DNA extraction using the DNeasy blood and tissue Kit (Qiagen)

To extract DNA from bacteria, a lysis step was added to the extraction protocol described by the manufacturer: a 1μL loopful of bacterial colonies was emulsified in 180 μL of lysis buffer containing 20 mM Tris-HCl, pH8, 2 mM EDTA, 1.2% Triton X-100, 20 mg/mL lysozyme, in a DNase/RNase free 1.5 ml Eppendorf tube and incubated in a heating block at 37°C for 1 hour, with mixing every 20 min. A DNase/RNase free 1.5 ml Eppendorf tube containing 180 μL of a home-made lysis buffer but no bacterial colonies was included as a NTC. Then, the manufacturer protocol, modified slightly by us, was followed. In brief, 25 μL of proteinase K and 200 μL of AL buffer were added to the preparation, vortexed for 15 sec and incubated in a heating block at 56°C for 30 min. The preparation was then vortexed for *ca*. 30 sec and incubated in a heating block at 72°C for 10 min. At the end of the incubation, 200 μL of ethanol at −20°C were added to the tube and vortexed for 15 sec, and the supernatant was transferred into a DNeasy columns and centrifuged for 1 min at 4500 g. Five hundred microliters of AW1 buffer were added to the column, which was then centrifuged for 1 min at 4500 g. This step was repeated after adding AW2 buffer with a centrifugation of 3 min at 6700 g. After each centrifugation, the collecting tube was discarded and replaced by a new one, except for the last step in which the collecting tube was replaced by a DNA/DNase/RNase-free 1.5 mL Eppendorf tube. Then, 100 μL of AE buffer, preheated to ca. 55°C, were carefully added to the column and then centrifuged for 1 min at 4500 g. The eluate was recovered, added to the top of the same column and centrifuged again for 1 min at 4500 g. Finally, the column was discarded and the eluate was kept at +5°C.

To extract DNA from swab samples, swab tips were placed into DNA/DNase/RNase-free 1.5 ml Eppendorf tubes containing nuclease-free water (Promega). The swab shafts were cut with a pair of sterile dissection scissors to allow closing of the tubes. Tubes were vortexed for about 5 mins and swabs removed using sterile forceps. Then, the tubes were centrifuged for 5 min at 8000 g. The supernatants were discarded and 150 μL of home-made lysis buffer, described above, was added to each pellet. This suspension was incubated at 37°C in a heating block for 1 hour. Then, the same procedure as described above for bacteria was followed. At the end of the extraction, the tube was incubated for 10 min at 95°C in a heating block to inactivate pathogens which could be contained in the samples.

To extract DNA from tissue samples, we proceeded in the same way as indicated above for the boiling method from tissue samples until obtaining a homogeneous suspension, using the home-made lysis buffer instead of nuclease-free water. Then, this suspension was incubated at 37°C in a heating block for about 1 hour, and the same procedure as described above for bacteria was performed. A NTC was included in the above procedures. This NTC consisted of a DNA/DNase/RNAse-free 1.5 mL Eppendorf tube, which followed the same treatment as clinical specimens, but in which there was no clinical specimen material.

### Primers and probes

The primers and probes used to detect the *tox* gene and *rpoB* genes for *C. diphtheriae* and *C. ulcerans/C. pseudotuberculosis* species identification were as described by De Zoysa *et al*. (21). For this study we introduced a conserved fragment of the 16S rRNA gene instead of the fragment of *gfp* gene (28) as the internal process control (IPC). The two primers and probe (u-16S) used to detect the 16S rRNA gene were designed with the software LC probe design2 (Roche). 16S rRNA sequences of known pathogenic or commensal species of the respiratory tract were aligned and a final selection of primers and probe was accomplished according to their universality (**Figure S1**). The sequences of primers and probes are given in **Table 2**.

**Table 2.**
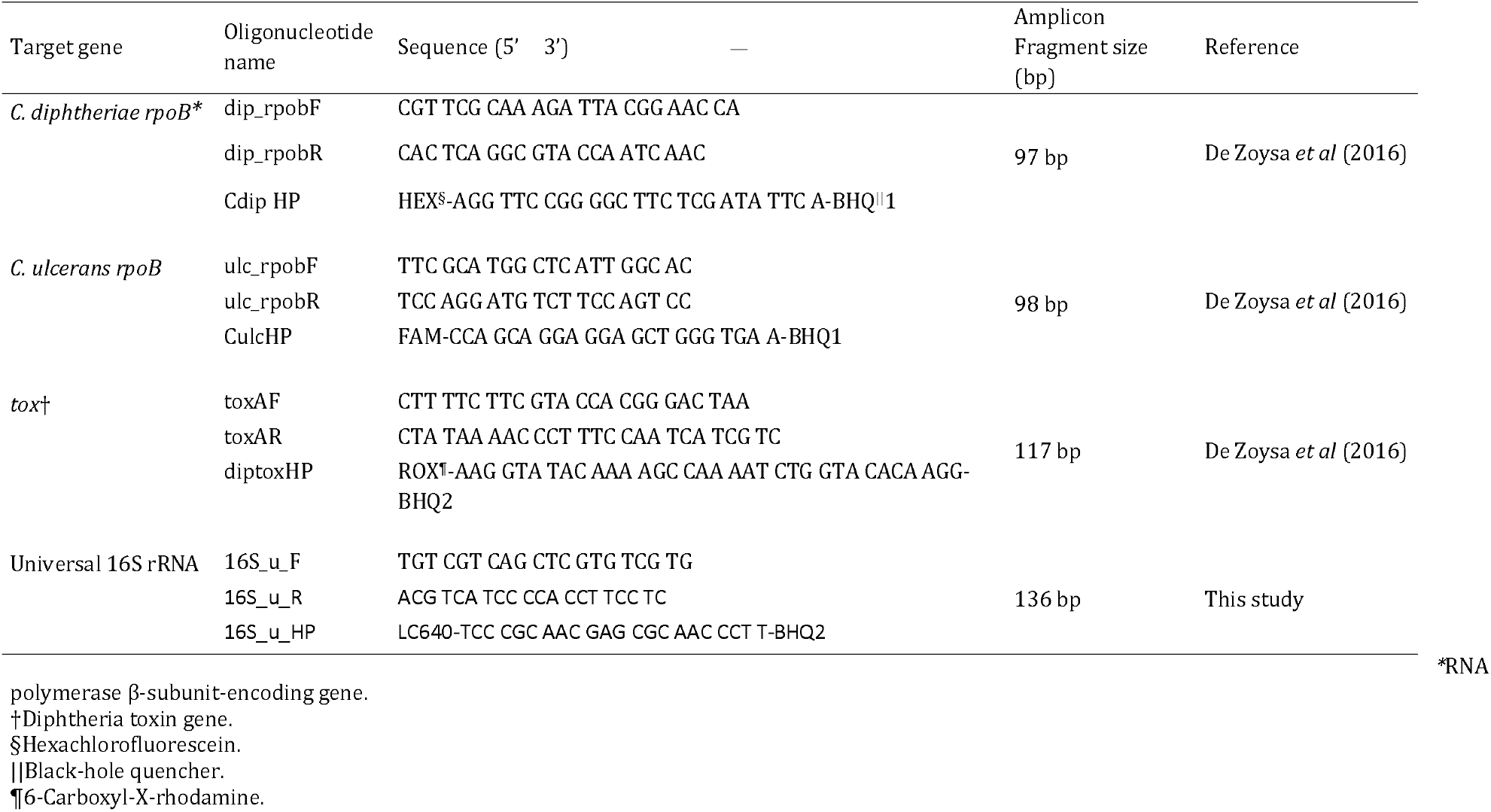
Oligonucleotides sequences and expected amplicon sizes of the four gene targets

### Reference end-point PCR method for *tox* gene detection

To detect the diphtheria *tox* gene, we used the conventional end-point PCR method described by Hauser *et al*. (20) modified by us to detect in parallel the bacterial 16S rRNA. In brief, DNA was extracted using the DNeasy Blood and tissue Kit (Qiagen) as described above. Two μL of DNA suspension were used in the final reaction described below. The PCR reaction was performed in a 50 μL volume containing: 0.25 μL of *Taq* DNA polymerase (5U/μL; Cat. No. 18038-026, Invitrogen, USA), 5 μL of 10X buffer (included in the *Taq* DNA polymerase kit), 2 μL of MgCl_2_ (50mM, included in the *Taq* DNA polymerase kit), 5 μL of 10 μM DT1 and DT2 primers (20), 1.25 μL of U5 and U4a primers (**Table S1**), 10 μL of deoxynucleoside triphosphates (2 mM, Cat. No. R1121, ThermoScientific, Lithuania). Thermocycling was performed on a AB 2720 thermocycler (Applied BioSystems, Singapore) with 1 cycle at 94°C for 3 min, followed by 35 cycles at 94°C for 20 s, 68 °C for 30 s, and 72 °C for 30 s and a final temperature of 15°C. The amplified products were resolved by electrophoresis on 3□% (w/v) agarose gels and visualized by ethidium bromide staining.

### Elek test for toxin production

Clinical isolates were tested for toxin production using Elek’s test modified by Engler *et al*. (15).

### Multiplex end-point PCR for species identification

A conventional multiplex end-point PCR was used to identify the isolates. This is a home-made end-point PCR adapted from these described by Pacheco *et al*. 2007(29) and Pimenta *et al*. 2008 (30). Briefly, DNA was extracted using the DNeasy Blood and tissue Kit, Qiagen as described above. Two μL of DNA suspension were used in the final reaction described below. The PCR reaction was performed in a 50 μl volume containing: 0.25 μL of *Taq* DNA polymerase (5U/μL, Cat. No. 18038-026, Invitrogen, USA), 5 μL of 10X Buffer (included in the *Taq* DNA polymerase kit), 2 μL of MgCl2 (50mM, included in the *Taq* DNA polymerase kit), 1 μL of each primers (10 μM) (29, 30), 5 μL of deoxynucleoside triphosphates (2 mM), (ThermoScientific, Cat. No. R1121, Lithuania). Thermocycling was performed on a thermocycler MJ Mini (BIO-RAD, Mexico) using 1 cycle at 95°C for 5 min, and 40 cycles at 95°C for 1 min, 58°C for 40 s, and 72°C for 1 min 30 s. Finally, the temperature was set to 72°C for 7 min and then at 15°C. The amplified products were resolved by electrophoresis on 3□% (w/v) agarose gels and visualized by ethidium bromide staining.

### qPCR

PCR assays were performed at the French National Reference Center, except where it is explicitly stated that they were performed at Public Health England. For qPCR amplification, we used the Qiagen Rotor-Gene Q (RGQ) thermocycler method as described by De Zoysa *et al*. (21). Some experiments were performed in parallel on a Roche LightCycler 480 II (LC480) thermocycler. Reaction mixture volumes were 20 μL in both thermocyclers. Each reaction mix comprised 10 μL of 2x Rotor-Gene Multiplex PCR Master Mix (Rotor-Gene Multiplex PCR Kit, catalogue no. 204774; Qiagen), 1 μL of a mix of primers and probes (to give final concentrations of 0.5 mM each primer and 0.2 mM each probe), 4 μL of H2O PCR grade and 5 μL of DNA template or H2O PCR grade. Five brands of H_2_O PCR grade were tested: Nuclease-free water (Cat. No. P119C, Promega, USA); UltraPure™ DNase/RNase-Free distilled water (Cat. No. 10977-035, Invitrogen™, UK); RNase-free water (included in the Rotor-Gene Multiplex PCR Kit, catalogue no. 204774, Qiagen, Germany), nuclease-free water (Cat. No. AM9937, Ambion, USA); and H_2_O PCR grade (included in the Kit LightCycler^®^ 480 Probes Master, Cat. No. 04707494001, Roche, Germany).

The cycling conditions were identical for both thermocyclers: an initial activation at 95°C for 5 min, followed by 45 cycles of denaturation at 95°C for 10 s followed by the hybridization/extension step at 60°C for 20 s. Acquisition of the fluorescence signal was set at 60°C during each cycle. The data analysis software used were Q-Rex (Qiagen) and LightCycler480 SW 1.5. For the determination of the cycle thresholds (Ct) value on the RGQ, the analysis options used were “Basic”, for all analyses, and “Slope correction” and/or “Take off Adjustment” if curves needed to be corrected. On the LC480 the second derivative method developed by Roche was used. Non-specific fluorescence from the HEX channel (*C. diphtheriae* target) can appear in the ROX channel (*tox* target) because the wavelengths of the two dyes are very close to each other (**Table S2**). To avoid this problem, the crosstalk compensation settings on the analysis options of the RGQ were used to define the channels that had to be compensated. Similarly, for the LC480, a colour compensation was performed to adjust the fluorescence results of each channel (**Table S3**). In the validation experiments at Public Health England, the PCRs were performed on an RGQ machine. When compared to the equivalent PCR using *gfp* as the IPC, the *gfp* reagents previously described (26) were used.

### Analytical sensitivity assays

The lower limits of detection (LLOD) of the qPCR assay were determined for each target at the French National Reference Center by using series of 10-fold dilutions of *C. diphtheriae* NCTC103356, *C. diphtheriae* NCTC10648, *C. ulcerans* NCTC12077 and *C. pseudotuberculosis* CIP102968^T^ DNAs at the initial concentration of 10 pg/μL. The online calculator page of Andrew Staroscik (https://cels.uri.edu/gsc/cndna.html) was used to calculate the number of genome copies corresponding to the DNA quantity. In the validation experiments at Public Health England, sensitivity of the qPCR assay was compared when using the u-16S IPC and the *gfp* IPC (26) using 2-fold serial dilutions of *C. diphtheriae* NCTC10648 and *C. ulcerans* NCTC12077 DNA between 40 and 5 genome copies/μL.

### Experimental robustness assays

To test the robustness of the method to temperature variation, we increased and decreased the temperatures of denaturation and annealing/elongation steps in the PCR program by 1°C, 2°C or 3°C. To test the effect of pipetting volume variation, we increased or decreased by 20% the volume of all PCR mix reagents simultaneously, while keeping fixed the volume of DNA template at 5 μL.

## Results

### Validation of u-16S primers and probe

A pair of primers and a probe that were maximally conserved on an alignment of 16S rRNA sequences (**Figure S1**) were defined (**Table 2**) and named the u-16S target. To test the newly-designed u-16S primers and probe for use as an appropriate control for bacterial DNA presence, we compared fluorescence signals obtained on the LC640 channel (used as dye for the u-16S target) using either DNA from bacteria or no template controls (NTCs). DNA at 10 pg/μL from four reference strains of the *diphtheriae* complex (NCTC10356, NCTC10648, NCTC12077 and CIP102968^T^) (**Table 1**) was tested on both the RGQ and LC480 thermocyclers, initially in simplex PCR. Crossing thresholds (Ct) were recorded in experiments in both instruments (although called crossing point, CP in the Roche system, we will call them Ct here for consistency). Fluorescence signals observed with bacterial DNA always had Ct values <27, whereas fluorescence signals from NTCs always showed Ct values ≥27 or higher (**Figure 1A**). This amplification signal was not expected for NTCs, and we suspected a contamination of the PCR grade H_2_O used, but it was observed systematically, even when using different brands and batches of PCR grade H2O. We conclude that the signal is presumably due to the presence of some residual genomic bacterial DNA in the qPCR mix reagents (31). The qPCR assay result on the LC640 channel was thus considered negative for the NTCs if the Ct value was ≥ 27, and was considered positive if the Ct value was ≤26.

**Figure 1.**
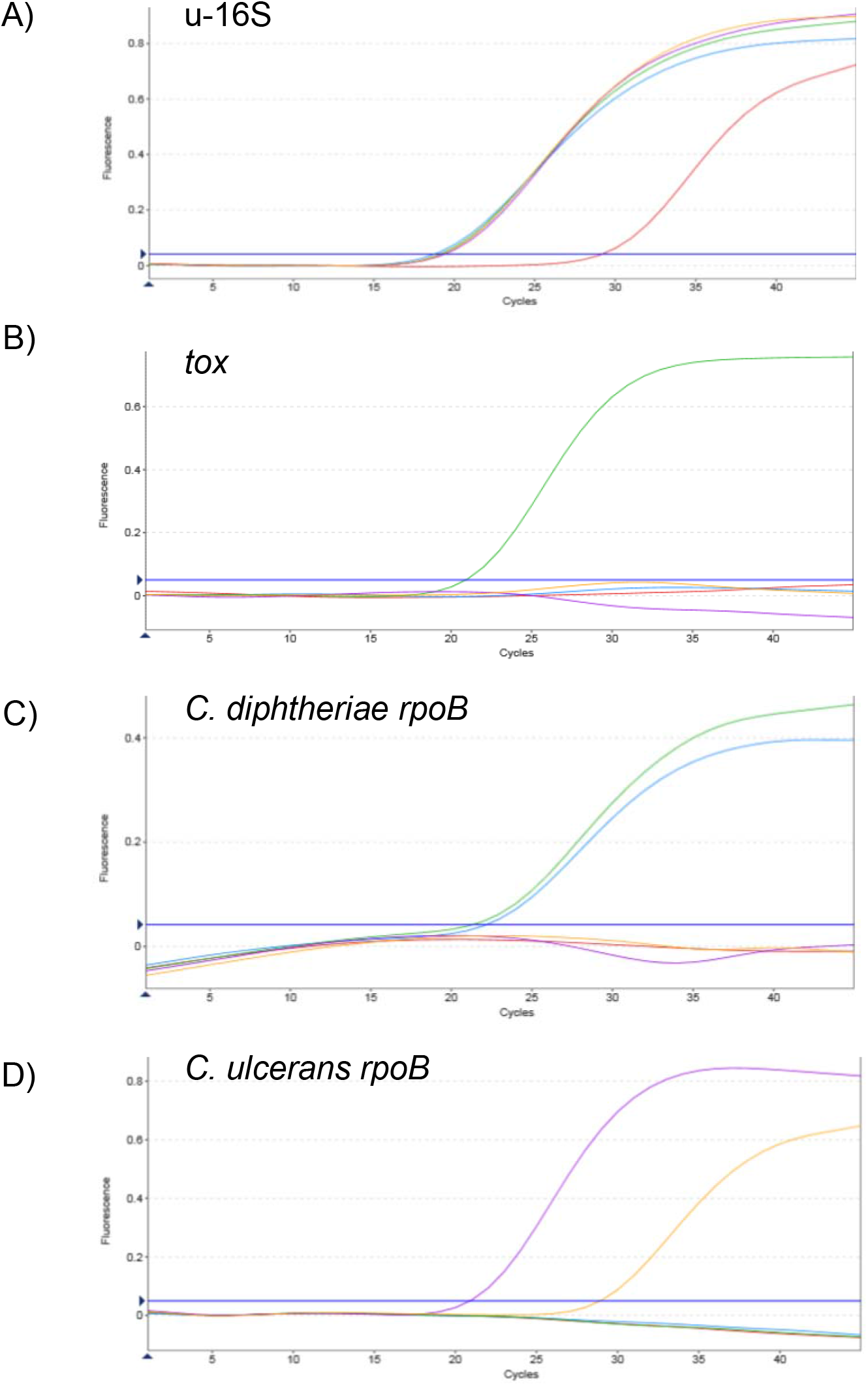
Example of qPCR curves for each of the targets.

We then tested whether the newly designed u-16S target signal interfered with the amplification signals expected in the channels HEX (*C. diphtheriae*), FAM (*C. ulcerans/C. pseudotuberculosis*) and ROX (*tox*) when used in quadruplex (4plex). We observed that fluorescence signals detected in the three channels were as expected for each target (**Figure 1, panels B-D**). Furthermore, no fluorescence signals in FAM, HEX and ROX channels were detected for the NTCs. Expected amplification of all targets was observed both on the RGQ and the LC480 platforms.

### Analytical sensitivity

The LLOD for *C. diphtheriae rpoB, C. ulcerans rpoB* and *tox* targets was 1 fg per μL, which corresponds to 0.37 genome copies per μL, or 1.86 genome copies per 5μL reaction. For *C. pseudotuberculosis*, the *rpoB* limit of detection was 186 genome copies per reaction. The LLOD obtained with *C. pseudotuberculosis* showed a lower sensitivity with the *C. ulcerans/C. pseudotuberculosis rpoB* target. Identical LLOD values were obtained on both thermocyclers. Regarding the u-16S target, between the dilutions 10 fg/μL and 0.1 fg/μL the Ct values were ca. 29 on the RGQ and *ca*. 33 on the LC480. As qPCR reagents contain DNA traces, it was not possible to observe the extinction of the fluorescence signal and therefore no LLOD could be determined for the u-16S target.

### Comparison of the two thermocyclers

We observed amplification curves in both thermocyclers for all targets tested. As mentioned above, LLOD were the same for all targets on both platforms. We noted that Ct values obtained with the two thermocyclers were slightly different (**Table 3**), but this did not impact the qualitative interpretation of the qPCR assay in terms of positive or negative results. We conclude that performance of the quadruplex qPCR assay on the two platforms was equivalent. For practical reasons, the subsequent experiments were performed only on the RGQ.

**Table 3.**
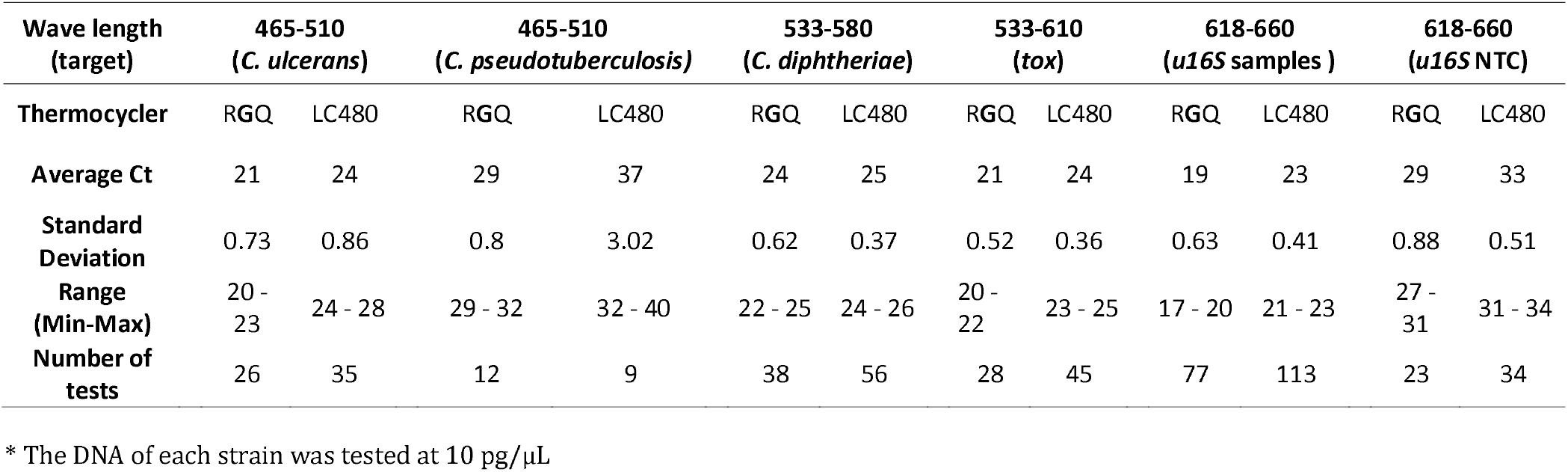
Comparison of crossing thresholds (Ct) values obtained using the Rotor-Gene Q (RGQ, Qiagen) and Lightcycler 480 II (LC480, Roche)^*^

### Analyses of strains, clinical isolates, and specimens

A panel of 43 bacterial DNA extracts from clinical isolates and strains belonging or not to the *C. diphtheriae* complex, and 16 clinical specimens, were analysed. This sample included 11 tox-positive isolates, among which six were non-toxigenic toxin bearing (NTTB) isolates. Fluorescence signals specific for *C. diphtheriae, C. ulcerans/C. pseudotuberculosis* and *tox* were always observed according to expectations, as defined using the conventional end-point PCR (**Table 1**). NTTB isolates were also positive for *tox* gene detection by the 4plex qPCR. These results confirm that the *tox* and species identification targets previously developed are correctly detected even in the presence of the novel u-16S target within the 4plex assay. In addition, fluorescence signals were detected for the u-16S target for all bacterial DNA extracts tested (all with Ct values ≤26), whether or not they were in the *C. diphtheriae* complex. This confirmed that the negative fluorescence signals in the channels HEX, FAM and ROX with non-*C. diphtheriae* complex isolates were not due to the accidental absence of bacterial DNA.

### Comparison of two DNA extraction methods

As amplifiable DNA is much faster to prepare using the boiling method (approximately 20 minutes) than using the kit extraction method (approximately 2 hours), we evaluated the boiling method as a template DNA preparation method for the 4plex qPCR. The 54 samples (isolates and clinical specimens) processed using this method were all positive for the u-16S channel (**Table 1**), showing that amplifiable DNA was obtained in all cases. Furthermore, samples processed using the boiling method were positive for all targets according to expectations based on the kit extraction method. We conclude that even though the DNA concentration is lower than with the kit extraction method, the boiling method can replace the kit extraction method for DNA preparation for the 4plex qPCR.

### Robustness

When increasing or decreasing the temperature of the thermocycler cycles by 3°C, Ct values did not vary by more than 2 cycles and no difference was observed in the interpretation of the qPCR amplification results (**Fig. S2**). The variation of the reagent volumes by +20% or −20% also had limited impact on the slopes and Ct values (< 2) compared to normal conditions (data not shown). These tests show that the 4-plex PCR is robust in the face of changes in experimental conditions.

### External validation of the qPCR

The modified qPCR using the u-16S IPC (instead of the *gfp* IPC) was validated in a second laboratory, the Respiratory and Vaccine Preventable Bacteria Reference Unit at Public Health England (RVPBRU-PHE), to confirm its portability and test its performance in comparison to the original method. Purified DNA from the toxigenic *C. diphtheriae* strain NCTC10648 and the non-toxigenic *C. ulcerans* strain NCTC12077 were tested in both versions of the qPCR at concentrations of 40, 20, 10 and 5 genome copies/μL in parallel over 20 runs to assess any effect on analytical sensitivity. The results showed that the sensitivity of the PCRs against the *C. diphtheriae rpoB, C. ulcerans/C. pseudotuberculosis rpoB* and the *tox* genes were essentially unaffected by changing the IPC; the differences in mean Ct values generated by both versions of the assay for comparative samples were less than 1 cycle (**Supplementary Table S4**). Positive results with the u-16S reagents did not generate any false positives for the other three targets. Ct values in the u-16S channel were all ≥28 cycles for NTCs (**Supplementary Table S4**; actual range 28.80 – 30.11). As expected, Ct values for the *gfp* IPC were consistently between 30 and 32 cycles regardless of the presence of target DNA.

## DISCUSSION

The quadruplex real-time PCR assay developed by De Zoysa *et al*. (26) for the identification of potentially toxigenic corynebacteria was an important advance in our diagnostic armamentarium. This includes an IPC consisting of a *gfp* gene target present in control DNA that is added to every PCR reaction in order to detect PCR inhibition. However, this IPC cannot distinguish between the analysis of a species that is not *C. diphtheriae, C. ulcerans* or *C. pseudotuberculosis* and a false negative due to the accidental lack of bacterial target DNA. However, in the theoretical case where the DNA extracted from a clinical sample was erroneously not added into the PCR mix, a positive signal will still be detected in the *gfp* channel. Negative results for *rpoB* and *tox* targets may lead to the wrong interpretation that no genetic material of *C. diphtheriae* complex was present.

Here we introduced a target corresponding to a universal fragment of the bacterial 16S rRNA gene instead of the *gfp* gene. This provides the ability to confirm the presence of bacterial DNA in the sample tubes in addition to the absence of PCR inhibition. Because it covers a broad range of bacteria, the fragment of u-16S is expected to be amplified if any bacterial DNA was introduced in the sample. The interpretation of the absence of signal for the u-16S target is that no bacterial DNA was present, or that the PCR amplification was inhibited, thus invalidating the assay.

We did detect some false positive signals on the LC640 (u-16S) channel when the NTCs were analysed. The cause of this is probably due to some residual genomic bacterial DNA present in the qPCR mix reagents. Contamination of the *Taq* DNA polymerase may originate from its production from bacterial cultures (31). As the Ct of these signals were always ≥ 27, whereas the Ct values from isolates or clinical samples were always <27 (typically between 17 and 20), we propose to treat 27 cycles as the background level in the u-16S channel and that any runs in which Ct values for the NTCs are ≥ 27 are valid.

We found a complete concordance of the improved 4-plex method regarding analyses of bacterial isolates as compared with the reference method, consistent with the results reported previously (26). In addition, we demonstrated that the qPCR can be used to detect the targets directly from clinical samples including pharyngeal swabs or pseudomembrane tissues. This is important because faster results can be obtained by avoiding the microbial culture step, which typically takes 18-24 hours. We also demonstrated that the improved 4-plex qPCR can be performed using DNA extracted using the boiling method from clinical samples or isolates, and that the method is robust within an important range of experimental variation of reagents volumes and thermocycler temperature drift that is unexpected to be exceeded in most laboratories. Moreover, the portability and performance of the modified 4plex qPCR were validated on an RGQ apparatus at RVPBR-PHE. Remarkably, the differences in mean Ct values between the two methods were less than 1 cycle in that laboratory. Finally, the LLOD defined using both thermocyclers were identical, providing flexibility to users in the choice of the thermocycler.

The use of a single target for *C. diphtheriae* and *C. belfantii* on the one hand, and of *C. ulcerans* and *C. pseudotuberculosis* on the other hand, does not allow for species discrimination within these pairs. *C. belfantii* can be identified by biotyping or by sequencing approaches (1). In humans, *C. pseudotuberculosis* is extremely rare and associated with contacts with goats or other production animals, whereas *C. ulcerans* is much more common. Therefore, positivity of *C. ulcerans/C. pseudotuberculosis* target assay may be interpreted in most cases as *C. ulcerans*. These two species are reliably distinguished using MALDI-TOF (17).

In conclusion, the improved 4-plex PCR method has the biological and technical characteristics required for the diagnostic of toxin gene-bearing strains of the *C. diphtheriae* species complex and we therefore recommend its deployment in medical biology and reference laboratories.

## Previous meeting presentations

The information in this work was not previously presented in any meeting.

## Acknowledgements

We acknowledge the help of Melody Dazas and Annick Carmi-Leroy for the microbiological characterization of the isolates.

