## Supplementary material for "Optimization and validation of a quadruplex real-time PCR assay for the diagnosis of diphtheria"

Figure S1

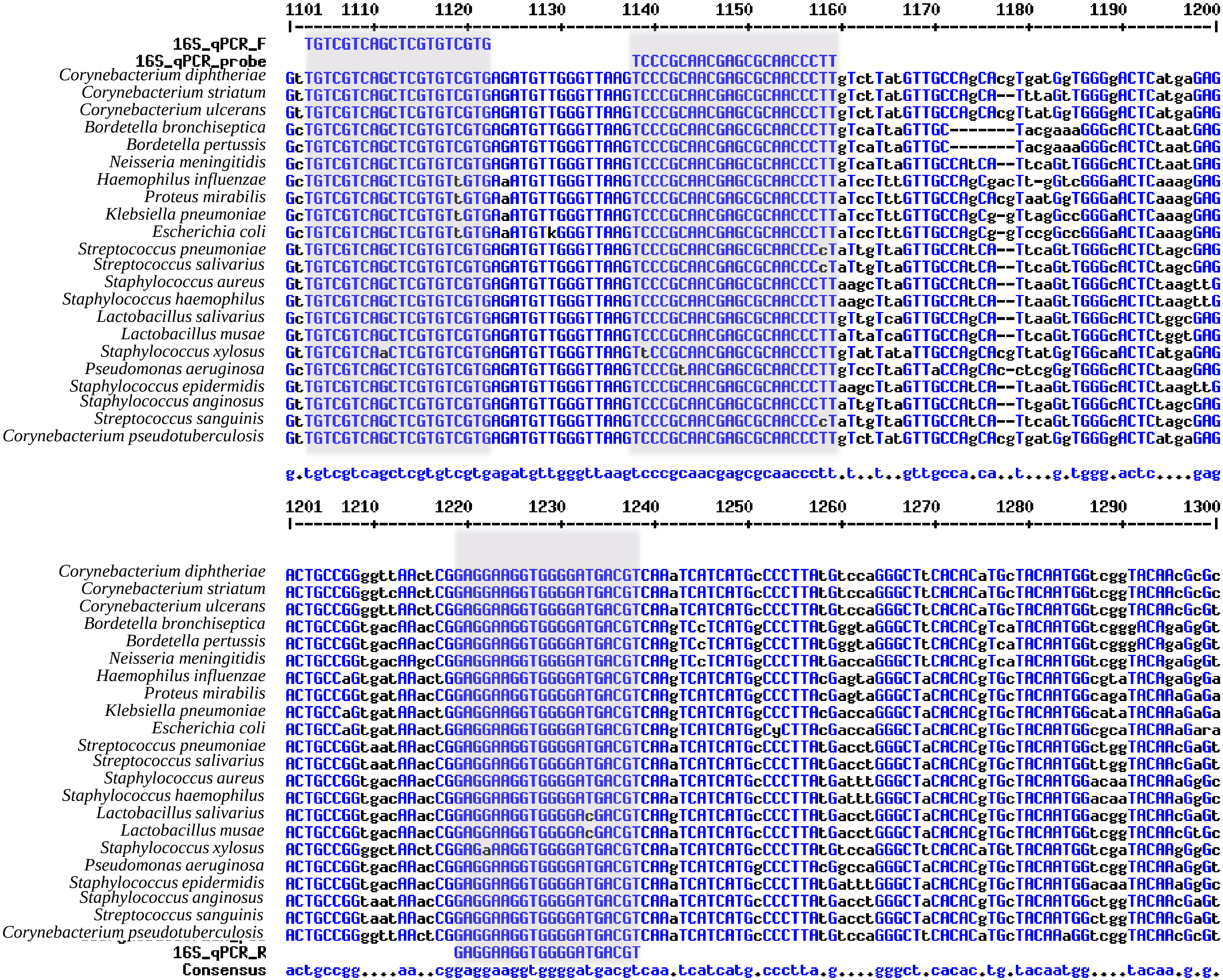

1101 1110 1120 1130 1140 1150 1160 1170 1180 1190 1200
16S_qPCR_F
16S_qPCR_probe
Corynebacterium diphtheriae
Corynebacterium striatum
Corynebacterium ulcerans
Bordetella bronchiseptica
Bordetella pertussis
Neisseria meningitidis
Haemophilus influenzae
Proteus mirabilis
Klebsiella pneumoniae
Escherichia coli
Streptococcus pneumoniae
Streptococcus salivarius
Staphylococcus aureus
Staphylococcus haemophilus
Lactobacillus salivarius
Lactobacillus musae
Staphylococcus xylosus
Pseudomonas aeruginosa
Staphylococcus epidermidis
Staphylococcus anginosus
Streptococcus sanguinis
Corynebacterium pseudotuberculosis
1201 1210 1220 1230 1240 1250 1260 1270 1280 1290 1300
16S_qPCR_R
Consensus

**Figure S1**. Alignment of 16S rRNA sequences in selected pathogens and their consensus sequence (below). The grey blocks from left to right correspond to forward primer, probe, and reverse primer, respectively.

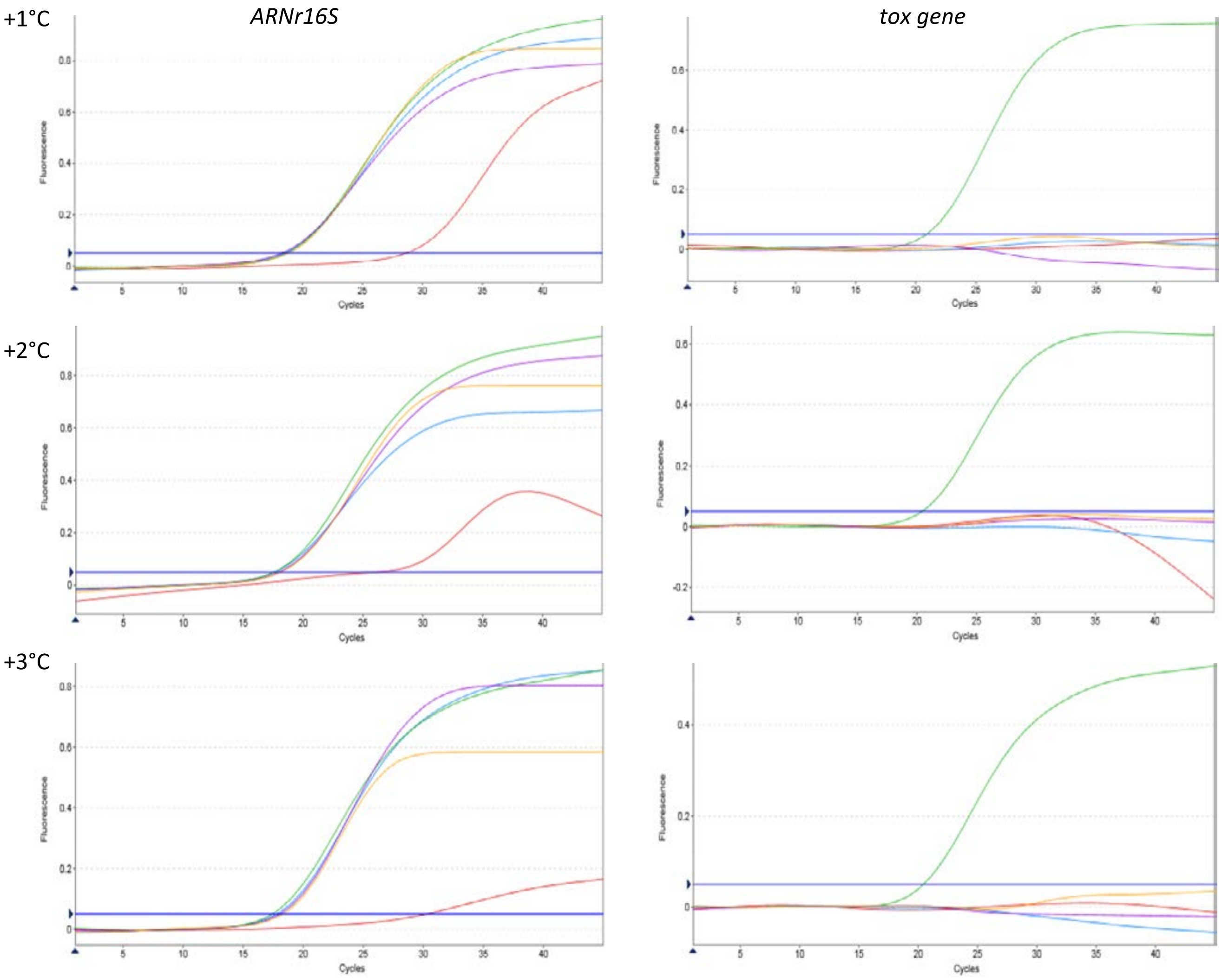

| Sample | $H_2O$ | | *C. diphteriae tox -* | | *C. diphteriae tox +* | | *C. ulcerans* | | *C. pseudotuberculosis* | |
|---|---|---|---|---|---|---|---|---|---|---|
| Temperature / Target | *ARN16S* | *tox* gene | *ARN16S* | *tox* gene | *ARN16S* | *tox* gene | *ARN16S* | *tox* gene | *ARN16S* | *tox* gene |
| No variation | 29 | > 45 | 19 | > 45 | 19 | 21 | 19 | > 45 | 19 | > 45 |
| + 1°C | 29 | > 45 | 19 | > 45 | 19 | 20 | 18 | > 45 | 19 | > 45 |
| + 2°C | 26 | > 45 | 18 | > 45 | 18 | 20 | 18 | > 45 | 18 | > 45 |
| + 3°C | 30 | > 45 | 18 | > 45 | 17 | 21 | 18 | > 45 | 18 | > 45 |

**Figure S2. Temperature robustness of 4plex qPCR method.**
The effect of the temperature variation on the qPCR was tested. For each amplification step the temperature was augmented from 1°C to 3°C. The upper panel shows the amplification curves, whereas the lower panel shows obtained Ct values of the u-*16S* and *tox* gene targets. No significant modifications of Ct were observed even at + 3°C, compared to the normal conditions. These results show the strong robustness of qPCR technique. Equivalent results were obtained  by diminishing the temperature from 1°C until 3°C (not shown).

**Table S1. Oligonucleotides sequences and expected amplicon sizes of the two gene targets, used in end-point PCR**

| Target gene | Oligonucleotide name | Sequence (5' - 3') | Amplicon fragment size (bp) | Reference |
|---|---|---|---|---|
| *tox* # | DT1 | CGG GGA TGG TGC TTC GCG | 910 | Hauser *et al*. (1993) |
| | DT2 | CGC GAT TGG AAG CGG GGT | | |
| 16S rRNA & | U5 | TCA AAG GAA TTG ACG GGG GC | 480 | This study |
| | U4a | AGG CCC GGG AAC GTA TTC A | | |

### Diphtheria toxin gene

& 16S rRNA gene

**Table S2**. **Color compensation method used on the LC480 II***

| Number of cycles | Steps | Temperature (°C) | Duration | Ramp rate | Analysis mode | Acquisition |
|---|---|---|---|---|---|---|
| 1 | Initial activation | 95 | 5 min | | | |
| 45 | Denaturation | 95 | 10 s | | | |
| | Annealing / Elongation | 60 | 20 s | | | |
| 1 | Temperature gradient | 95 | 10 s | 4.4 | | |
| | | 40 | 30 s | 2 | | |
| | | 80 | | 0.05 | Continuous | 3°C/s |
| | | 40 | 30 s | 2.2 | | |

* The combination of filters was: FAM (excitation: 465 nm - emission: 510 nm), HEX (533-580), ROX (533-610) and LC640 (618-660).

Compensation parameters were: Melting factor: 1; Quanti factor: 10; and Max integration time: 2 seconds

**Table S3. Comparison of analytical sensitivity of the multiplex PCR against *C. diphtheriae rpoB*, *C. ulcerans/C. pseudotuberculosis rpoB* and the *tox* genes when using a) the *gfp* IPC described by de Zoysa *et al.*, 2016 and b) the u-16S IPC described in this study.**

| **Gene target** | **Target DNA (gc/µl) #** | **a) Assay with *gfp* IPC** | | **b) Assay with u-16S IPC** | |
|---|---|---|---|---|---|
| | | **Mean Ct** | **Standard deviation** | **Mean Ct** | **Standard deviation** |
| *C. diphtheriae rpoB* | *C. diphtheriae* NCTC10648: | | | | |
| | 40 | | | | |
| | 20 | 31.68 | 0.45 | 31.34 | 0.44 |
| | 10 | 32.60 | 0.42 | 32.25 | 0.65 |
| | 5 | 33.53 | 0.90 | 33.04 | 0.68 |
| | | 34.56 | 1.17 | 34.57 | 1.13 |
| *C. ulcerans/* | *C. ulcerans* NCTC12077: | | | | |
| *C. pseudotuberculosis* | 40 | | | | |
| *rpoB* | 20 | 29.00 | 0.48 | 28.97 | 0.49 |
| | 10 | 30.36 | 0.54 | 30.16 | 0.52 |
| | 5 | 31.58 | 1.08 | 31.64 | 0.84 |
| | | 33.90 | 1.32 | 34.38 | 2.04 |
| *tox* gene | *C. diphtheriae* NCTC10648: | | | | |
| | 40 | | | | |
| | 20 | 30.79 | 0.34 | 30.73 | 0.42 |
| | 10 | 31.83 | 0.42 | 31.75 | 0.75 |
| | 5 | 32.78 | 1.01 | 33.13 | 1.80 |
| | | 33.79 | 0.83 | 34.55 | 2.64 |
| Internal process | *C. diphtheriae* NCTC10648: | | | | |
| control | 40 | | | | |
| | 20 | 31.34 | 0.75 | 27.95 | 0.51 |
| | 10 | 31.47 | 0.76 | 28.63 | 0.42 |
| | 5 | 31.37 | 0.65 | 29.23 | 0.51 |
| | | 31.47 | 0.80 | 29.75 | 0.45 |
| | *C. ulcerans* NCTC12077: | | | | |
| | 40 | | | | |
| | 20 | | | | |
| | 10 | 31.23 | 0.72 | 28.41 | 0.36 |
| | 5 | 31.35 | 0.73 | 29.01 | 0.53 |
| | | 31.39 | 0.73 | 29.34 | 0.46 |
| | NTC | 31.38 | 0.79 | 29.73 | 0.35 |
| | | 31.42 | 0.77 | 29.34 | 0.45 |

### gc: genome copies. Purified DNA from the toxigenic *C. diphtheriae* strain NCTC10648 and the non-toxigenic *C. ulcerans* strain NCTC12077 were tested in both versions of the qPCR at concentrations of 40, 20, 10 and 5 genome copies/microliter in parallel over 20 runs.

**Table S4. qPCR detection channels.**

| | | | Rotor-Gene Q | | Roche LC480 II | |
|---|---|---|---|---|---|---|
| **Probe target** | **Dye/Quencher combination** | **Channel** | **Excitation (nm)** | **Detection (nm)** | **Excitation (nm)** | **Detection (nm)** |
| *C. ulcerans/*<br>*C. pseudotuberculosis* | FAM/BHQ-1 | Green | 470±10 | 510±5 | 465 | 510 |
| *C. diphtheriae* | HEX/BHQ-1 | Yellow | 530±5 | 557±5 | 533 | 580 |
| Diphtheria toxin gene | Rox/BHQ-2 | Orange | 585±5 | 610±5 | 533 | 610 |
| Universal 16S rRNA gene | LC640/BHQ-2 | Red | 625±5 | 660±10 | 498 | 640 |
